# Effects of the cannabinoid receptor agonist CP-55,940 on incentive salience attribution

**DOI:** 10.1101/767103

**Authors:** Ali Gheidi, Lora M. Cope, Christopher J. Fitzpatrick, Benjamin N. Froehlich, Rachel Atkinson, Coltrane K. Groves, Clair N. Barcelo, Jonathan D. Morrow

**Affiliations:** Department of Psychiatry, University of Michigan; Addiction Center, University of Michigan; Neuroscience Graduate Program, University of Michigan; Molecular and Behavioral Neuroscience Institute, University of Michigan

## Abstract

Pavlovian conditioned approach paradigms are used to characterize the nature of motivational behaviors in response to stimuli as either directed toward the cue (i.e., sign-tracking) or the site of reward delivery (i.e., goal-tracking). Recent evidence has shown that activity of the endocannabinoid system increases dopaminergic activity in the mesocorticolimbic system, and other studies have shown that sign-tracking behaviors are dependent on dopamine. Therefore, we hypothesized that administration of a cannabinoid agonist would increase sign-tracking and decrease goal-tracking behaviors. Forty-seven adult male Sprague Dawley rats were given a low, medium, or high dose of the cannabinoid agonist CP-55,940 (*N*=12 per group) or saline (*N*=11) before Pavlovian conditioned approach training. A separate group of rats (*N*=32) were sacrificed after PCA training for measurement of cannabinoid receptor type 1 (CB1) and fatty acid amide hydrolase (FAAH) using *in situ* hybridization. Contrary to our initial hypothesis, CP-55,940 dose-dependently decreased sign-tracking and increased goal-tracking behavior. CB1 expression was higher in sign-trackers compared to goal-trackers in the prelimbic cortex, but there were no significant differences in CB1 or FAAH expression in the infralimbic cortex, dCA1, dCA3, dorsal dentate gyrus, or amygdala. These results demonstrate that cannabinoid signaling can specifically influence behavioral biases toward sign- or goal-tracking. Pre-existing differences in CB1 expression patterns, particularly in the prelimbic cortex, could contribute to individual differences in the tendency to attribute incentive salience to reward cues.

## Introduction

The incentive sensitization theory of addiction focuses on the role played by cues in the development of compulsive drug-seeking, drug-taking, and relapse (1, 2). Cues are ubiquitous in our environment and can serve a predictive function, adaptively identifying stimuli as rewarding or harmful. Cues can also become attractive and desired in their own right when they acquire incentive motivational properties or *incentive salience* through Pavlovian processes, sometimes leading to maladaptive behaviors such as addiction.

Pavlovian conditioned approach (PCA) paradigms are used to characterize motivational behaviors in response to cues as directed either toward the cue itself (i.e., sign-tracking) or the site of reward delivery (i.e., goal-tracking). Briefly, one version of this paradigm involves placing rats into a chamber with a lever and a food magazine, with the appearance of the lever (conditioned stimulus) reliably predicting the delivery of a food pellet (unconditioned stimulus) into the food magazine. Over many trials, individual animals develop one of three motivational response patterns: sign-tracking, goal-tracking, or intermediate responding. Sign-tracking rats reliably approach and engage with the lever (e.g., pressing and biting it) whenever it is available, indicating the lever has been imbued with incentive salience, whereas goal-tracking rats glance at the lever and then approach and engage with the food magazine prior to food delivery. Intermediate responders exhibit both types of behaviors. Importantly, all groups learn the predictive value of the lever, but only for sign-trackers does the lever acquire incentive salience. This paradigm is relevant for understanding addiction processes because individual animals that are identified as sign-trackers show faster acquisition of drug self-administration (3), increased preference for drugs over food (4), and greater reinstatement of drug-seeking than goal-trackers (5–7).

Previous work has shown that dopamine is necessary for sign-tracking but not goal-tracking (8), consistent with the known involvement of dopamine in limbic structures such as the amygdala, hippocampus, and prefrontal cortex in incentive salience attribution (9, 10). The endocannabinoid system, including the enzyme fatty acid amide hydrolase (FAAH), has been implicated as a modulator of mesocorticolimbic dopamine transmission (11). FAAH regulates endocannabinoid levels by specifically degrading N-arachidonoylethanolamine (anandamide), thereby decreasing the concentration of endocannabinoids that bind to cannabinoid receptors such as CB1 (12). Importantly, there are individual differences in levels of both endocannabinoids and FAAH in the mesocorticolimbic system of both rats and humans (13, 14). These findings suggest potential relevance of endocannabinoids and FAAH in sign- and goal-tracking behaviors specifically and in addiction processes more broadly (12), but few studies have explored this possibility directly.

To this end, we studied rats using a combination of behavioral (i.e., PCA) and molecular genetic methods (e.g., *in situ* hybridization) to assess the role of cannabinoids and FAAH in sign- and goal-tracking behaviors. In Experiment 1, rats underwent PCA training after being injected with the cannabinoid agonist CP-55,940 or saline. We hypothesized that CP-55,490 would increase dopamine transmission, thereby increasing sign-tracking behaviors and decreasing goal-tracking behaviors in a dose-dependent manner. In Experiment 2, we measured baseline levels of FAAH and CB1 mRNA in goal-trackers, sign-trackers, and intermediate responders in order to determine whether individual differences in FAAH expression is associated with PCA behavioral phenotypes. We hypothesized that sign-tracking behavior would correlate with lower levels of FAAH and higher levels of CB1 mRNA, especially in limbic areas known to participate in reward learning.

## Materials and Methods

### Experiment 1

#### Subjects

Adult male Sprague Dawley rats (*N*=47) were pair-housed on a 12:12 light/dark cycle and had *ad libitum* access to food and water throughout the experiments. This study was approved and conducted in accordance with the guidelines set out by the Institutional Animal Care and Use Committee of the University of Michigan.

#### Pavlovian conditioned approach and the cannabinoid agonist CP-55,940

Pavlovian conditioning was conducted in modular conditioning chambers (24.1 cm width × 20.5 cm depth × 29.2 cm height; MED Associates, Inc.; St. Albans, VT) contained within individually sound-attenuated cabinets equipped with ventilation fans to provide ambient white noise. A red house light was located on one wall of each chamber. The opposite wall contained a centrally placed pellet magazine, and an illuminated, retractable lever counterbalanced on the left or right of the pellet magazine. When activated, the retractable lever extended into the chamber and was illuminated by an LED light within the lever housing. A pellet dispenser delivered banana-flavored food pellets (45 mg; Bioserv; Frenchtown, NJ) into the pellet magazine. Head entries into the pellet magazine were detected by an infrared sensor.

For the first 2 days, animals underwent one 30-minute pre-training session per day during which each rat was placed in a PCA cage with the house light on, and 50 food pellets were delivered on a variable-time (VT) 30-second schedule. The following day, rats were injected (1 mL/kg; i.p.) with either the cannabinoid agonist CP-55,940 (*n*=36; C112-10MG; Sigma-Aldrich, Inc.) or a control vehicle (*n*=11; 5% ethanol, 5% Cremphor EL [C5135-500G; Sigma-Aldrich, Inc.], and 90% saline) 30 minutes before testing. Rats given CP-55,940 received a concentration of 10 ug/kg (*n*=12), 50 ug/kg (*n*=12), or 100 ug/kg (*n*=12). All animals underwent PCA training for seven days. Sessions were about 40 minutes in length and consisted of 25 trials on a VT 90-second schedule. For each trial, the illuminated lever was extended into the cage for 8 seconds, followed immediately by delivery of one food pellet into the magazine. We henceforth refer to this 7-day period of Experiment 1 as the *acquisition phase*. Based on the robust behavioral effects observed, the highest dose was chosen for a drug crossover whereby rats that were previously injected with high-dose CP-55,940 (i.e., 100ug/kg) were injected with the control vehicle, and rats that were previously injected with the control vehicle were injected with high-dose CP-55,940. The rats then underwent four additional days of PCA sessions. We henceforth refer to the sessions conducted after the switch in drug administration as the *crossover phase*.

#### Statistical Analysis

We used a previously developed PCA index formula (15) to generate PCA scores ranging from −1 to 1, with a score of −1 corresponding to strong goal-tracking behavior, a score of +1 corresponding to strong sign-tracking behavior, and a score of 0 corresponding to intermediate behavior with no bias toward the lever or food magazine. PCA index scores were calculated by averaging measures of response bias, latency, and probability: [([lever presses - magazine entries] / [lever presses + magazine entries]) + ([magazine entry latency - lever press latency] / 8) + (lever press probability - magazine entry probability)] / 3. In addition to PCA index, the number, latency, and probability of lever presses and magazine entries were used as dependent measures.

For the purposes of statistical analysis, we discarded the first three days of PCA testing during which animals learned to perform the task. We conducted linear mixed modeling in SAS v9.4 using PROC MIXED. Four models were tested: 1) the effects of group (control vehicle, low dose, medium dose, or high dose) and day (days 4, 5, 6, and 7) (acquisition phase); 2) the effects of group (control vehicle vs. all drug groups combined) (acquisition phase); 3) the effects of treatment (control vehicle vs. high dose), period (pre-crossover vs. post-crossover), and the interaction between treatment and period (crossover design); and 4) the effects of treatment (control vehicle vs. high dose), day (days 4, 5, 6, 7, 8, 9, 10, and 11), and the interaction between treatment and day (crossover design). For model 4, we were specifically interested in the comparison of each of the post-crossover days (i.e., days 8, 9, 10, and 11) with the final precrossover day (i.e., day 7).

### Experiment 2

#### Subjects

Adult male Sprague Dawley rats (*N*=32) were pair-housed on a 12:12 light/dark cycle and had *ad libitum* access to food and water throughout the experiments. This study was approved and conducted in accordance with the guidelines set out by the Institutional Animal Care and Use Committee of the University of Michigan.

#### Pavlovian conditioned approach

All rats experienced a two-day pre-training procedure in which they were placed in the PCA cages while the house light remained on and the lever remained retracted as 50 food pellets were delivered on a VT 30-second schedule. Following pre-training, the rats underwent 5 days of testing sessions on the PCA paradigm described in Experiment 1. PCA index scores were computed, and rats were classified as goal-trackers (i.e., −1 to -.5), intermediate responders (i.e., -.5 to +.5), or sign-trackers (i.e., +.5 to +1) (16). After the last testing session, all rats were sacrificed and their brains flash frozen for radiolabeled ^35^S *in situ* hybridization. From these animals, we extracted the brains of 8 sign-trackers, 8 goal-trackers, and 8 intermediate responders for sectioning. These 24 subjects were selected based on PCA index scores that best represented the full distribution (−1 to +1).

#### Tissue processing

One week after PCA training rats were decapitated, and the brains were rapidly removed, frozen in isopentane (−75°C), and then stored at −80°C until further processing. Coronal brain sections (10 μm) were cut on a cryostat in 12 serial sections (at 170-μm intervals) throughout the prefrontal cortex, hippocampus, and amygdala, and thaw-mounted onto Superfrost/Plus slides (Fisher Scientific, Pittsburgh, PA, USA). Slides were stored at −80°C until *in situ* hybridization was performed.

### Molecular Cloning

#### FAAH

The cDNA FAAH insert was donated in plasmid form by Dr. Ken Mackie at the University of Indiana. The insert was then removed from the parent vector (PCDNA 3) and ligated into restriction sites of pBluscript. (This was done to eliminate the use of SP6 over T3 for a superior polymerase vector.) The parent vector was inserted into JM109 cells and grown with established bacterial transformation protocols. The end product was sent for sanger sequencing to the University of Michigan Sequencing Core. The plasmid was then linearized to both sense and anti-sense strands and later conjugated to S^35^ for *in situ* hybridization.

#### CB1

Rat cerebellar RNA was extracted and converted to cDNA to be used as a template for polymerase chain reaction (PCR). The PCR product was then ligated into the pSC-A-amp/kan vector and transformed into StrataClone competent cells. The plasmid including the insert was then sent to the University of Michigan Sequencing Core for sanger sequencing in order to confirm presence and orientation of the insert. Upon confirmation, the plasmid was linearized with BamHI and HindIII and made into sense and antisense for S^35^ riboprobe labeling.

#### Riboprobe labeling

FAAH and CB1 DNA linearized for antisense was mixed with ^35^S-conjugated to uracil and adenine (Perkin Elmer) incubated at 37°C for 1.5 hours. 1 uL of DNAse was added to reaction, mixed, and incubated at room temperature for 15 minutes. The labeled riboprobe was separated from free ^35^S via a Biorad Micro Bio-Spin 6 Chromatography column. Gel was resuspended via inversion of column, and flow-through was discarded. Column was spun at 1,000g for 2 minutes before volume of labeling reaction was brought to 75 uL with water. The column was placed in an Eppendorf tube and spun at 1,000g for 4 minutes. After mixing, 1 uL of flowthrough was counted. 1 uL of dithiothreitol (DTT) was added, and the labeled probe was stored at −80°C until ready for use.

#### Radiolabeled *in situ* hybridization (ISH) with S^35^

ISH was performed as previously described (17). Briefly, slides were removed from −80°C and placed immediately in 4% buffered formaldehyde at room temperature for 1 hour. Slides were then washed three times in 2X SSC for 5 minutes per each wash. Slides were placed in .1M triethanolamine of pH 8.0 with .25% vol/vol acetic anhydride and incubated at room temperature for 10 minutes before being washed in water. Sections were dehydrated in graded alcohols from 50% to 100% and allowed to air dry. Radioactive probe was then diluted in hydbridization buffer (50% formamide, 10% dextran sulfate, 3X SSC, 50 mM sodium phosphate, pH 7.4, 1X denhardts, and .1 mg/mL yeast tRNA) to give between 1 and 2 million dpms/35 to 70 uL. DTT was added to achieve a final concentration of 10 mM. 35uL to 70 uL of radioactive probe and hybridization buffer mixture was placed on slides via a coverslip, and slides were placed in a hybridization box with 50% formamide soaked filter paper in the bottom. Slides were incubated in the hybridization oven overnight at 55°C. The next day, coverslips were removed with 2X SSC and washed three times in 2X SSC for 5 minutes per each wash. Sections were placed in a 37°C RNAse A solution (10 mM Tris HCl, pH 8.0, .5 M NaCl, 200 ug/mL RNAse A) for 60 minutes. Sections were washed for 5 minutes each consecutively in 2X SSC, 1X SSC, and .5X SSC. Sections were placed in 65°C .1X SSC for 1 hour. Sections were rinsed in distilled water, dehydrated through graded alcohols from 50% to 100%, and allowed to air dry.

Sections were placed on Kodak Biomax MR film (Eastman Kodak, Rochester, NY, USA). Sections were exposed for one week. The specificity of the hybridization signal was confirmed by control experiments using sense and anti-sense probes (Supplemental Figure S1). Autoradiograms were scanned via a Microtek ScanMaker 1000XL (Fontana, CA) and the scanner was driven by Lasersoft Imaging (SilverFast) software (Sarasota, FL). In addition, a macro was used which enabled signal to be automatically determined.

#### Quantification of radioactive signal

Regions of interest were the cingulate/prelimbic cortex/infralimbic cortex, dorsal/ventral CA1 (dCA1), dorsal/ventralCA3 (dCA3), dorsal/ventral dentate gyrus, and amygdala. Signals for each region of interest were calculated by using the mean optical density. A measurement of each tissue slice on the slide was taken. Duplicate slices and measurements for the left and right hemispheres for each region were averaged together to produce one measurement for each region per rat. Quantification was performed by experimenters blind to the conditions of the animals. Tissue slices with poorly defined or unclear regions of interest were discarded.

#### Statistical Analysis

Quantification of the optical density of tissue slices exposed to the FAAH/CB1 ^35^S-cDNA probe is a measure of the levels of mRNA expression in any given brain region, with optical density scores ranging from 0 (no FAAH/CB1) to 1 (maximal FAAH/CB1 levels). IBM SPSS statistics v25 was used to analyze CB1 and FAAH data. Independent samples t-tests were used to compare sign- and goal-trackers on CB1 and FAAH expression in each region. In addition, Pearson product-moment correlations were computed to examine associations between optical density measurements (i.e., FAAH and CB1 expression) and PCA measures (i.e., number of lever presses, lever press latency, number of magazine entries, magazine entry latency, and PCA index score). For all analyses, significance was set at .05.

## Results

### Experiment 1: CP-55,490

#### Model 1

This model tested the main effect of group (four levels) and main effect of day (days 4–7) during the acquisition phase (Figure 1). There was a significant main effect of group for lever press number, latency, and probability; magazine entry latency and probability; and PCA index (all *p*s < .023). There was also a trend toward an effect of group for magazine entry number (*p* = .063). For all dependent measures, the control vehicle group was significantly different from the high dose group (all *p*s < .048), with the high dose group having fewer lever presses, a longer lever press latency, a lower lever press probability, more magazine entries, a shorter magazine entry latency, a higher magazine entry probability, and a lower PCA index, all indicating greater goal-tracking. There was also a main effect of day for lever press number and latency (*p* = .002 and .009, respectively). See Table 1 for all Model 1 results.

**Figure 1.**
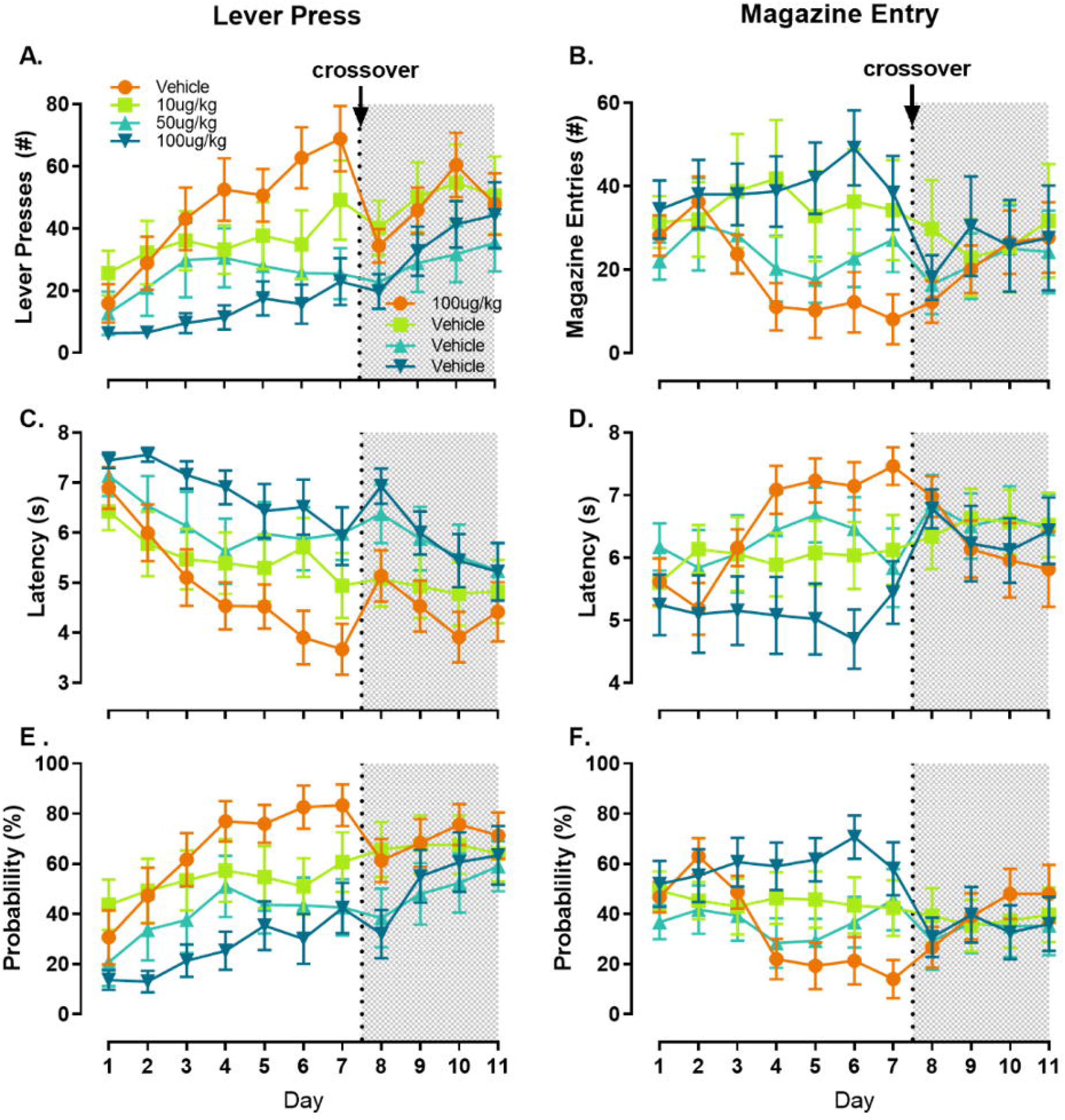
Pavlovian conditioned approach behavior in response to presentation of a retractable lever paired with food delivery. Sign-tracking behavior as measured by number (A), latency (C), or probability (E) of lever presses, as well as goal-tracking behavior as measured by number (B), latency (D), or probability (F) of food magazine entries plotted as a function of day of training. Colors and symbols denote the dose of CP-55,490 or vehicle that each experimental group was given 30 min prior to testing each day. The experiment followed a crossover design such that on day 8 the vehicle group was switched to 100 μg/kg PC-55,490, and all other groups were switched to vehicle. Data after the crossover are plotted in the shaded right-hand portion of the graphs. All data are presented as mean ± SEM of responses during the lever presentation.

**Table 1.** Experiment 1 (CP-55,490) Models 1 and 2 (acquisition phase). Model 1 tests the main effect of group (1, 2, 3, or 4) and the main effect of day. Main effect degrees of freedom (*df*) = 3, 138. Group comparison *df* = 138. Group comparisons are reported when the main effect is either significant at *p* < .05 or trending at *p* < .08. Model 2 tests the main effect of group (vehicle vs. drug). Degrees of freedom (*df*) = 1, 138. LP, lever press; ME, magazine entry; PCA, Pavlovian conditioned approach. †*p* < .08, \**p* < .05, \*\**p* < .01, \*\*\**p* < .001

| MODEL 1 | LP number | LP latency | LP probability | ME number | ME latency | ME probability | PCA index |
| --- | --- | --- | --- | --- | --- | --- | --- |
| Main Effect of Group | $F = 5.91, p < .001^{***}$ | $F = 4.63, p = .004^{**}$ | $F = 5.74, p = .001^{**}$ | $F = 2.49, p = .063^{\dagger}$ | $F = 3.31, p = .022^{*}$ | $F = 3.37, p = .020^{*}$ | $F = 6.44, p < .001^{***}$ |
| Main Effect of Day | $F = 5.02, p = .002^{**}$ | $F = 4.00, p = .009^{**}$ | $F = 1.90, p = .133$ | $F = 1.48, p = .222$ | $F = 0.56, p = .642$ | $F = 0.87, p = .461$ | $F = 2.32, p = .078^{\dagger}$ |
| Group Comparisons: |  |  |  |  |  |  |  |
| vehicle vs. low | $t = 2.16, p = .033^{*}$ | $t = -2.06, p = .041^{*}$ | $t = 2.30, p = .023^{*}$ | $t = -2.00, p = .048^{*}$ | $t = 1.67, p = .097$ | $t = -1.75, p = .082$ | $t = 2.43, p = .016^{*}$ |
| vehicle vs. medium | $t = 3.12, p = .002^{**}$ | $t = -2.78, p = .006^{**}$ | $t = 3.09, p = .002^{**}$ | $t = -0.85, p = .398$ | $t = 1.19, p = .234$ | $t = -1.05, p = .294$ | $t = 2.62, p = .010^{**}$ |
| vehicle vs. high | $t = 4.02, p < .001^{***}$ | $t = -3.56, p < .001^{***}$ | $t = 3.97, p < .001^{***}$ | $t = -2.46, p = .015^{*}$ | $t = 3.10, p = .002^{**}$ | $t = -3.08, p = .002^{**}$ | $t = 4.38, p < .001^{***}$ |
| low vs. medium | $t = 0.99, p = .325$ | $t = -0.74, p = .463$ | $t = 0.81, p = .418$ | $t = 1.18, p = .241$ | $t = -0.49, p = .626$ | $t = 0.71, p = .476$ | $t = 0.19, p = .850$ |
| low vs. high | $t = 1.91, p = .059^{\dagger}$ | $t = -1.54, p = .127$ | $t = 1.71, p = .089$ | $t = -0.47, p = .639$ | $t = 1.46, p = .147$ | $t = -1.36, p = .176$ | $t = 1.99, p = .049^{*}$ |
| medium vs. high | $t = 0.92, p = .360$ | $t = -0.80, p = .425$ | $t = 0.90, p = .370$ | $t = -1.65, p = .102$ | $t = 1.95, p = .053^{\dagger}$ | $t = -2.07, p = .040^{*}$ | $t = 1.80, p = .074^{\dagger}$ |
| Day Comparisons: |  |  |  |  |  |  |  |
| 4 vs. 5 | $t = -0.56, p = .574$ | $t = 0.38, p = .706$ | | | | | $t = -0.50, p = .616$ |
| 4 vs. 6 | $t = -1.04, p = .302$ | $t = 0.75, p = .457$ | | | | | $t = 0.55, p = .585$ |
| 4 vs. 7 | $t = -3.59, p < .001^{***}$ | $t = 3.14, p = .002^{**}$ | | | | | $t = -1.96, p = .052^{\dagger}$ |
| 5 vs. 6 | $t = -0.47, p = .638$ | $t = 0.37, p = .714$ | | | | | $t = 1.05, p = .295$ |
| 5 vs. 7 | $t = -3.02, p = .003^{**}$ | $t = 2.76, p = .007^{**}$ | | | | | $t = -1.46, p = .147$ |
| 6 vs. 7 | $t = -2.55, p = .012^{*}$ | $t = 2.39, p = .018^{*}$ | | | | | $t = -2.51, p = .013^{*}$ |
| MODEL 2 | LP number | LP latency | LP probability | ME number | ME latency | ME probability | PCA index |
| Main Effect of Group | $F = 13.61, p < .001^{***}$ | $F = 11.42, p < .001^{***}$ | $F = 13.99, p < .001^{***}$ | $F = 4.50, p = .036^{*}$ | $F = 5.55, p = .020^{*}$ | $F = 5.36, p = .022^{*}$ | $F = 13.66, p < .001^{***}$ |

#### Model 2

This model tested the main effect of group (two levels) during the acquisition phase (Figure 1). There was a significant main effect of group (control vehicle vs. all drug doses combined) for all seven dependent measures (all ps < .036), again with the drug group showing greater goal-tracking behaviors than the control vehicle group (e.g., fewer lever presses and more magazine entries). See Table 1 for all Model 2 results.

#### Model 3

This model tested the main effect of treatment (control vehicle vs. high dose), main effect of period (pre- vs. post-crossover), and interaction between treatment and period for the crossover design (Figure 1). There was a significant interaction between treatment and period for lever press number, lever press latency, lever press probability, and PCA index (all *p*s < .001). There was also a trend toward effects for magazine entry number and latency (*p* = .070 and .057, respectively). Both of the within-group interaction effects (i.e., control-vehicle/pre vs. high-dose/post and high-dose/pre vs. control-vehicle/post) were significant for lever press number, lever press probability, magazine entry number, magazine entry latency, and PCA index (all *p*s < .020). See Table 2 for all Model 3 results.

**Table 2.**
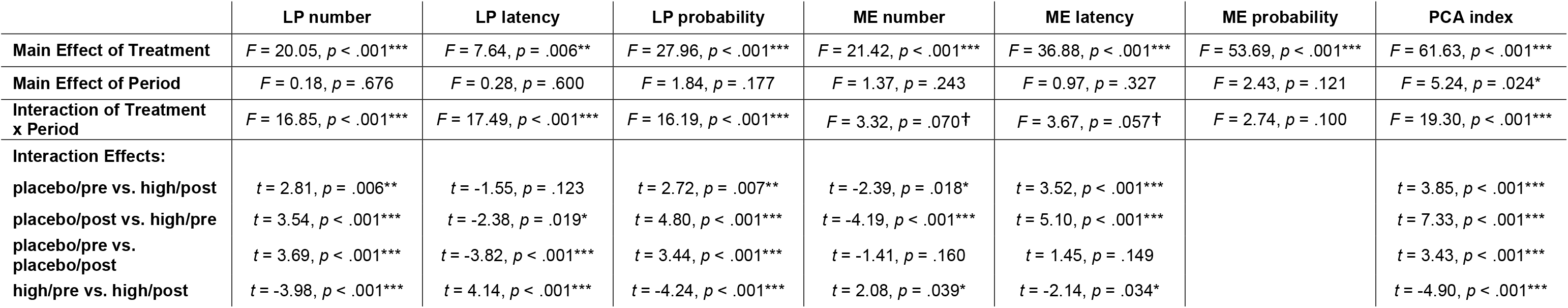
Experiment 1 (CP-55,490) Model 3 (crossover design), testing main effect of treatment, main effect of period, and interaction of treatment and period. Treatment is coded as 1 = vehicle, 2 = high dose; Period is coded as 1 = pre-crossover, 2 = post-crossover. Main effect and interaction degrees of freedom (*df*) = 1, 159. Interaction effects *df* = 159. Interaction effects are reported when the interaction is either significant at *p* < .05 or trending at *p* < .08. LP, lever press; ME, magazine entry; PCA, Pavlovian conditioned approach. †*p* < .08, \**p* < .05, \*\**p* < .01, \*\*\**p* < .001

#### Model 4

This model tested the main effect of day (days 4–11), main effect of treatment (control vehicle vs. high dose), and interaction between day and treatment for the crossover design (Figure 1). There was a significant interaction between day and treatment for lever press number, lever press latency, lever press probability, and PCA index (all *p*s < .005). There was a significant difference between day-7/control-vehicle vs. day-8/high-dose for lever press number, lever press latency, lever press probability, and PCA index (all *p*s < .040). There was also a significant difference for day-7/high-dose vs. day-8/control-vehicle for lever press latency (*p* = .019). See Table 3 for all Model 4 results.

**Table 3.**
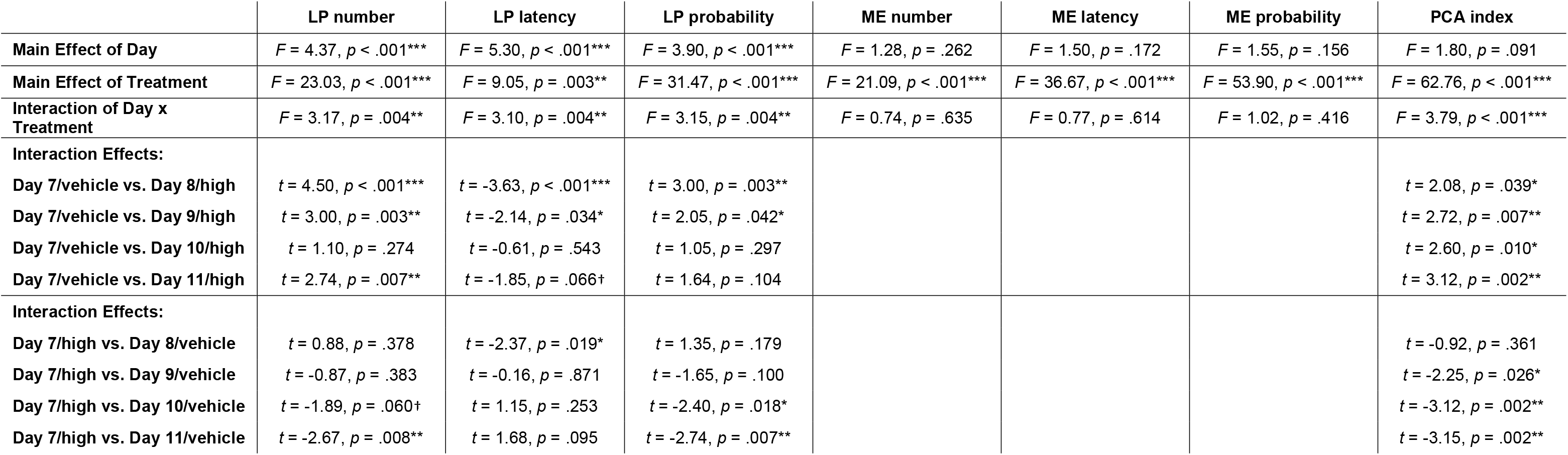
Experiment 1 (CP-55,490) Model 4 (crossover design), testing main effect of day, main effect of treatment, and interaction of day and treatment. Treatment is coded as 1 = vehicle, 2 = high dose. Main effect of day and interaction degrees of freedom (*df*) = 7, 147. Main effect of treatment *df* = 1, 147. Interaction effects *df* = 147. Interaction effects are reported when the interaction is either significant at *p* < .05 or trending at *p* < .08. LP, lever press; ME, magazine entry; PCA, Pavlovian conditioned approach. †*p* < .08, \**p* < .05, \*\**p* < .01, \*\*\**p* < .001

### Experiment 2: FAAH and CB1

CB1 and FAAH expression were quantified in the cingulate cortex, prelimbic and infralimbic prefrontal cortex, dorsal and ventral CA1 (dCA1), dorsal and ventralCA3 (dCA3), dorsal and ventral dentate gyrus, and amygdala (See Supplemental Figure S2 for regions of interest). CB1 expression was higher in sign-trackers compared to goal-trackers in the prelimbic cortex, t(14) = 2.58, *p* = .02 (Figure 2). There were no significant differences in CB1 or FAAH expression in the infralimbic cortex, dCA1, dCA3, dorsal dentate gyrus, or amygdala (all *p*s > .05). There was a significant correlation between PCA index score and CB1 expression in the prelimbic cortex, *r* = .48, *p* = .022. There were no other significant correlations between PCA index score and expression of either CB1 or FAAH (all *p*s > .05).

**Figure 2.**
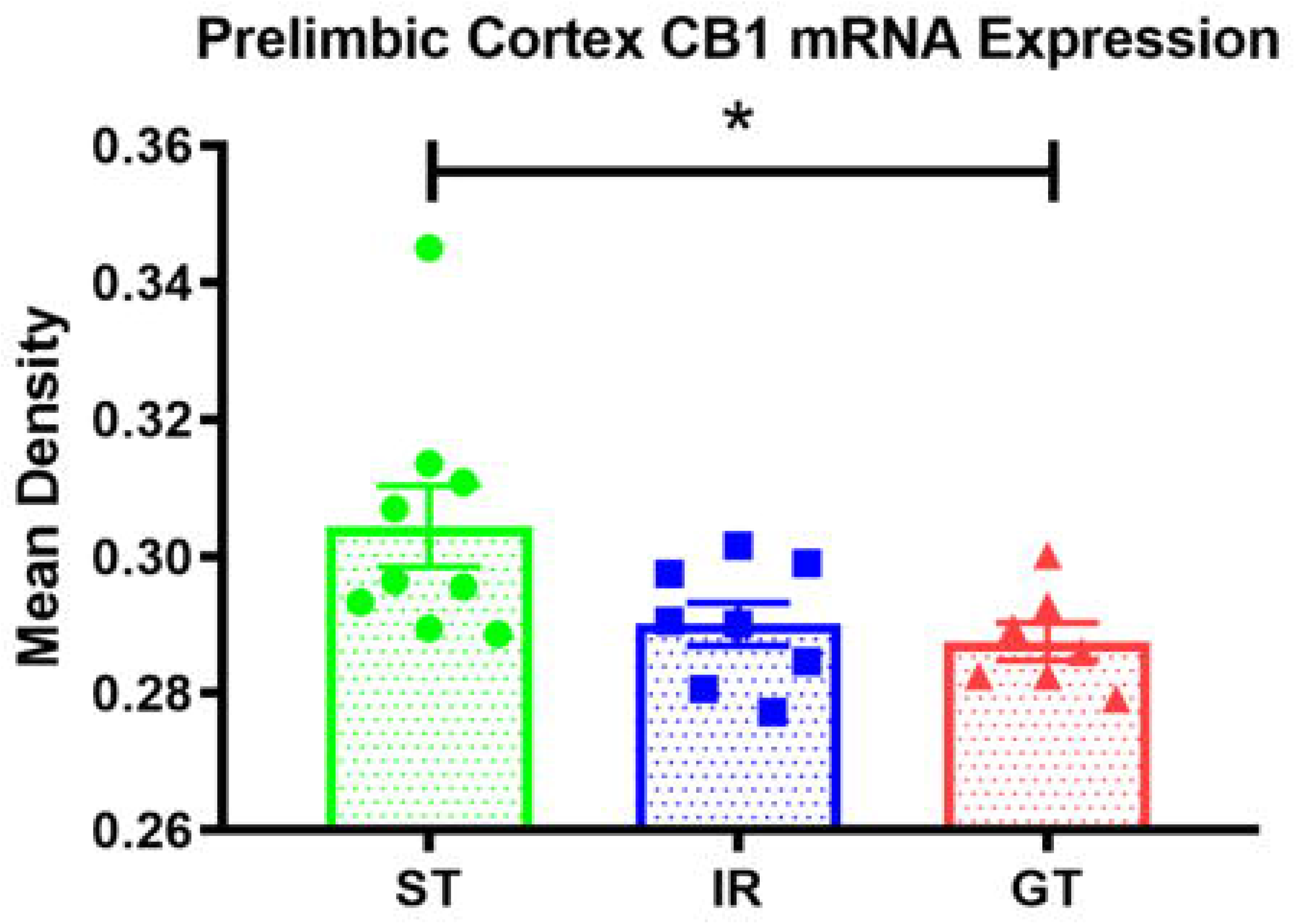
CB1 mRNA expression in the prelimbic cortex. The different colors represent behavioral phenotypes classified as sign-trackers (ST), goal-trackers (GT), or intermediate responders (IR). Bars represent mean ± SEM, and symbols are individual data points. * = *p* < .05 sign tracker vs. goal tracker.

## Discussion

These data demonstrate that systemic administration of the non-selective cannabinoid agonist CP-55,940 dose-dependently decreases sign-tracking and increases goal-tracking behavior. This effect is reflected in a number of different measures, including number of responses, latency of responses, and probability of responses. The fact that this drug increased one conditioned response while decreasing another conditioned response in the same subject makes it less likely that the observed effects were due to non-specific impairments of motor activity, cognition, or motivated behavior in general. The histological experiments did not detect differences in FAAH or CB1 expression in most of the brain regions quantified, but CB1 expression in the prelimbic cortex was higher in sign-trackers compared to goal-trackers.

The finding that cannabinoid agonism increases goal-tracking and decreases sign-tracking behavior is surprising and directly opposite of our original hypothesis. As noted in the introduction, dopaminergic activity in the nucleus accumbens generally promotes sign-tracking activity (8, 18–20), and cannabinoid agonists are known to disinhibit dopamine release into the nucleus accumbens (21–24). Furthermore, sign-tracking behavior predicts increased reinstatement of food and drug self-administration (6, 7, 25), and cannabinoid agonists also generally promote reinstatement of food- and drug-seeking behaviors (26–31). However, dopamine is thought to increase sign-tracking by attributing incentive salience properties specifically to cues that are associated with reward. Accordingly, dopaminergic, glutamatergic, and electrical activity within the nucleus accumbens is time-locked to the appearance of reward-cues whenever sign-tracking is observed (8, 32, 33). It may be that endogenous cannabinoids normally facilitate sign-tracking behavior, but use of a long-acting cannabinoid agonist disrupts the normal sign-tracking process by promoting dopamine release to extraneous events, thereby obscuring cue-specific salience signals. This interpretation is supported by reports that the cannabinoid inverse agonist rimonabant, instead of having the opposite effect of the agonist used in this study, also decreases sign-tracking behavior (34). Additionally, behavioral shifts away from sign-tracking similar to those observed in this study have been reported by other groups using amphetamine and dopamine agonists (35, 36).

Regardless of the explanation of the specific effects of CP-55,940 on sign- and goal-tracking, these data make it clear that cannabinoids can profoundly influence the acquisition and expression of PCA behavior. Because genetic polymorphisms that affect FAAH and CB1 expression in humans have been linked with vulnerability to addiction and other psychiatric disorders (37, 38), we hypothesized that variation in expression of these proteins might contribute to sign- vs. goal-tracking phenotypes. While we detected individual differences in FAAH expression, these differences were not correlated with sign- and goal-tracking behavior. This argues against an overwhelming influence of FAAH expression on PCA phenotypes, though it is important to mention that individual differences in PCA behaviors are most likely not due to a single neurobiological factor, but rather reflect a number of converging factors that together influence motivational behavior. It is possible that a relationship between FAAH levels and PCA phenotypes may be have been obscured by variation in other neural factors such as dopamine receptors, cholinergic activity, and other mechanisms that affect reward learning.

Despite such potential challenges, we did find higher levels of CB1 expression in the prelimbic prefrontal cortex of sign-trackers relative to goal-trackers, indicating that increased cannabinoid activity in the prelimbic may promote sign-tracking behavior. While this may appear to contrast with our pharmacological data, cannabinoids are known to exert effects on reward-seeking behaviors that are highly regionally specific. For example, increasing endogenous cannabinoid signaling by systemic inhibition of FAAH decreases alcohol consumption, but FAAH inhibition targeted to the prefrontal cortex increases alcohol consumption, and rats selectively bred to self-administer alcohol have decreased FAAH expression specifically in the prefrontal cortex (39, 40). Overall, these findings lend further support to the notion that pre-existing differences in regionally specific cannabinoid activity may contribute to behavioral differences between sign- and goal-trackers.

## Acknowledgements

The authors would like to thank Dr. Ken Mackie at the University of Indiana for his helpful advice and for providing the plasmids used for *in situ* hybridization experiments. We would also like to thank Dr. Stanley Watson at the University of Michigan for providing some material support for the *in situ* hybridization experiments.

