## Supplemental Information for "Effects of the cannabinoid receptor agonist CP-55,940 on incentive salience attribution"

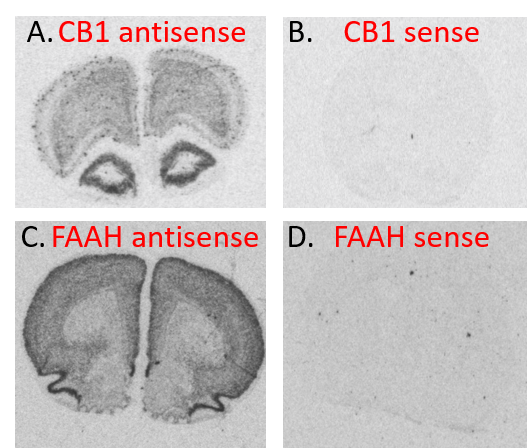

**Supplemental Figure S1.** Specificity of CB1 and FAAH mRNA S^35^ riboprobes. Dark S^35^ staining of antisense riboprobes is present in the prefrontal cortex for both CB1 (A) and FAAH (C), but absent for the sense riboprobes (B and D).

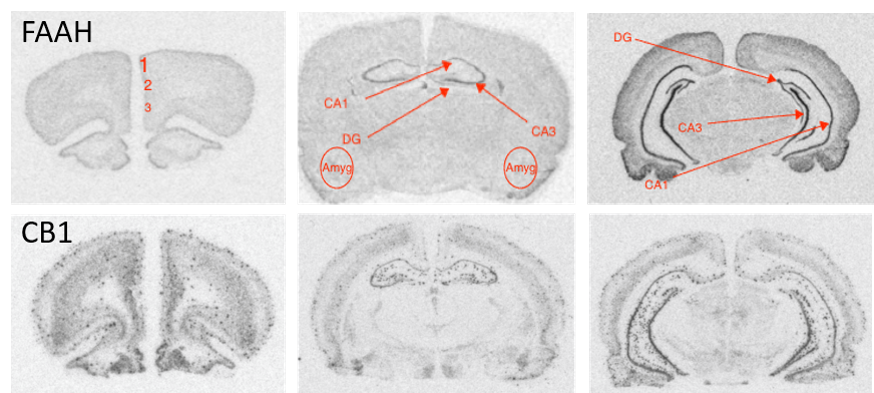

**Supplemental Figure S2.** Representative images of regions of interest after S^35^ radioactive mRNA labeling of FAAH (top) and CB1 (bottom). 1 = cingulate cortex; 2 = infralimbic cortex; 3 = prelimbic cortex; Amyg = amygdala; CA1 = hippocampal area CA1; CA3 = hippocampal area CA3; DG = dentate gyrus.
