## Supplementary figures and images for "Effects of the cannabinoid receptor agonist CP-55,940 on incentive salience attribution"

### Figure S1

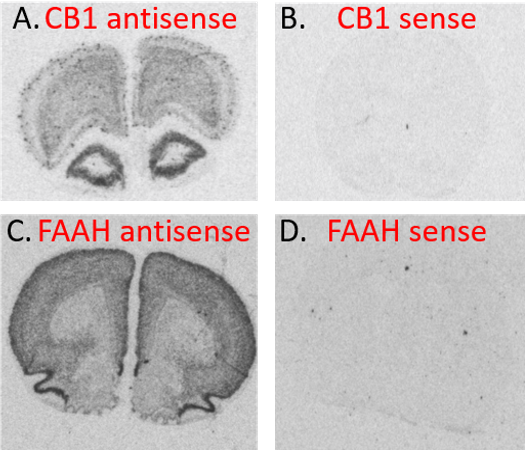

### Figure S2

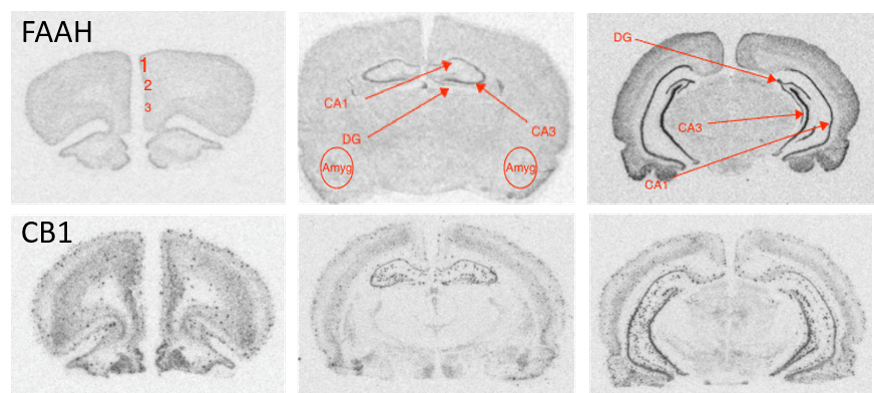
